# Phase separation provides a mechanism to reduce noise in cells

**DOI:** 10.1101/524231

**Authors:** F. Oltsch, A. Klosin, F. Jülicher, A. A. Hyman, C. Zechner

**Affiliations:** Max Planck Institute of Molecular Cell Biology and Genetics, 01307 Dresden, Germany; Center for Systems Biology Dresden, 01307 Dresden, Germany; Max Planck Institute for the Physics of Complex Systems, 01187, Dresden Germany

## Abstract

A central problem in cellular control is how cells cope with the inherent noise in gene expression. Although transcriptional and posttranscriptional feedback mechanisms can suppress noise, they are often slow, and cannot explain how cells buffer acute fluctuations. Here, by using a physical model that links fluctuations in protein concentration to the theory of phase separation, we show that liquid droplets can act as fast and effective buffers for gene expression noise. We confirm our theory experimentally using an engineered phase separating protein that forms liquid-like compartments in mammalian cells. These data suggest a novel role of phase separation in biological information processing.

## Main text

Stochasticity in gene expression causes substantial cell-to-cell variability in protein concentration, even in genetically identical cell populations grown in the same environmental conditions^1,2^. This stochasticity stems both from random binding and modification events in transcription and translation (*intrinsic* noise), as well as cell-to-cell differences in factors affecting gene expression, such as the local microenvironment^3^, ribosome abundance or ATP availability^1^ (*extrinsic* noise). Together, intrinsic and extrinsic noise can lead to cell to cell differences in protein concentrations over several orders of magnitude^4^. Yet, living organisms display an extraordinary degree of robustness and can exhibit precise spatial and temporal organization.

Liquid-liquid phase separation provides a potential mechanism to buffer concentration variability^8,13^. This is because in a phase separating system, the concentrations inside and outside the droplets are constrained by thermodynamic laws. When the total concentration changes, the droplets change in number and size, while the concentration outside of the droplets will remain largely unaffected. We illustrate this for the phase separating protein 2NT-DDX4^YFP^ (Fig. 1A and materials and methods)^*14*^which forms liquid droplets in low salt buffer in a concentration dependent manner (Fig. 1B). Once a threshold concentration is reached, the bulk phase concentration outside the droplets stays largely insensitive to changes in total protein concentration (Figs. 1B-D). This *in vitro* experiment shows that liquid compartments could serve as dynamic reservoirs that buffer cell-to-cell variability in protein concentrations (Fig. 1E).

**Figure 1.**
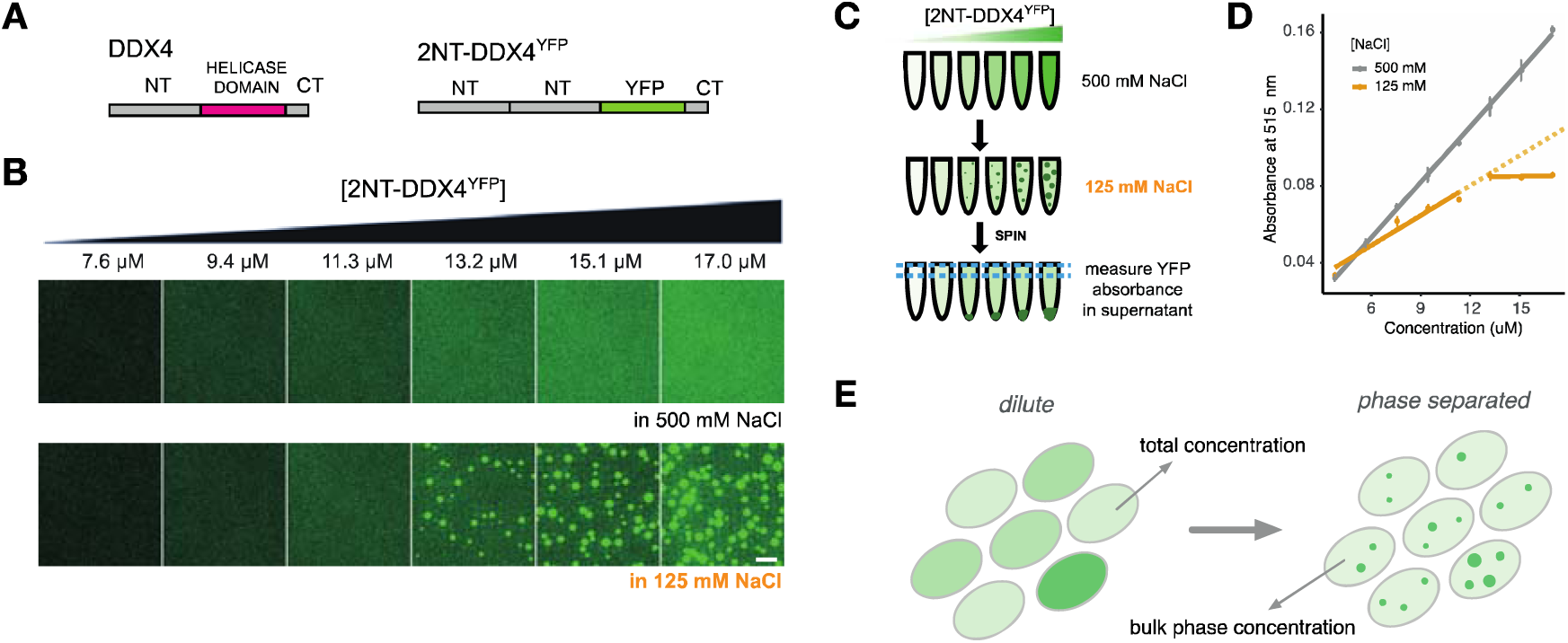
Phase separation buffers variations in protein concentration. **(A)** Diagram of the phase-separating model protein 2NT-DDX4^YFP^. The helicase domain of the human DDX4 protein was replaced with yellow fluorescent protein (YFP) and the disordered N-terminus (NT) was duplicated. **(B)** Purified 2NT-DDX4^YFP^ phase separates in a concentration-dependent manner. Scale bar = 10 µm. Images were set to high contrast to demonstrate the change of intensity in the dilute, bulk phase. **(C)** Purified and diluted 2NT-DDX4^YFP^ was triggered to phase-separate by reducing the salt concentration. The droplet phase was separated by centrifugation, the supernatant containing the dilute phase was recovered and quantified as a function of increasing protein concentration. **(D)** Quantification of the protein concentration in the recovered supernatant phase using spectrophotometry. Mean and standard deviation from 3 measurements at each concentration and in two salt conditions are plotted. Linear fits were performed separately for concentrations below and above the value where phase separation was observed by microscopy **(E)** Phase-separation could buffer variability in protein concentration (represented by color intensity) between different cells by forming liquid compartments of variable number and size.

To test the efficacy of phase separation in buffering gene expression noise, we developed a physical model based on the thermodynamics of liquid-liquid phase separation^11^. We consider a binary mixture consisting of a solvent and a phase separating protein (Fig. 2A and Supplementary text S.1.1). When the total protein concentration in the cell (which we denote by 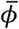) exceeds a certain threshold *ϕ*^*^, the mixture separates into a dilute bulk phase of concentration *ϕ*_+_ and a condensed droplet phase with concentration *ϕ*_−_. Importantly, in a mesoscopic droplet system these concentrations fluctuate with time, even when the total concentration 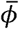 is fixed and the system is at equilibrium. This is because the exchange of protein material between droplet and bulk phase is subject to thermal noise, which may become significant in small compartments like cells. These fluctuations represent a lower limit to the concentration noise in phase separating systems. To characterize this limit, we calculated the equilibrium probability distribution of the bulk phase protein concentration associated with our thermodynamic model for different, but fixed total concentrations 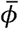 (Fig. 2B). Our analysis shows that when 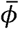 is above the threshold concentration *ϕ*^*^ (which we call the “buffering regime”), the mean bulk phase protein concentration remains close to the threshold protein concentration ((*ϕ*_+_) ≈ *ϕ*^*^, whereas (*ϕ*_+_) denotes the mean of (*ϕ*_+_), consistent with *in vitro* experiments (Fig. 1D). The corresponding noise strength in the bulk phase (which we define as the squared coefficient of variation CV[*ϕ*_+_]^2^ = σ[*ϕ*_+_]^2^/(*ϕ*_+_)^2^ with σ[*ϕ*_+_]^2^ = (*ϕ*_+_^2^) -(*ϕ*_+_)^2^ as the variance of *ϕ*_+_) decreases inversely with (*ϕ*_+_) and correspondingly also with *ϕ*^*^ (Fig. 2C). This has two important implications. First, the relationship between mean and noise strength of the bulk phase protein concentration is consistent with *Poissonian* noise. Poissonian noise is often considered the minimal achievable noise in protein expression^4,7^, because it arises when protein molecules are produced and turned over with constant rates, with no additional sources of stochasticity. Secondly, both the mean, 〈*ϕ*_+_〉 and the noise strength, CV[*ϕ*_+_]^2^ are effectively set by the threshold protein concentration, *ϕ*^*^, which is independent of the total protein concentration, 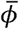. Taken together, this suggests that as long as a system phase separates, the noise in the bulk phase deviates only weakly from the *Poisson* limit, even if protein concentration exhibits fluctuations or cell-to-cell variability.

**Figure 2.**
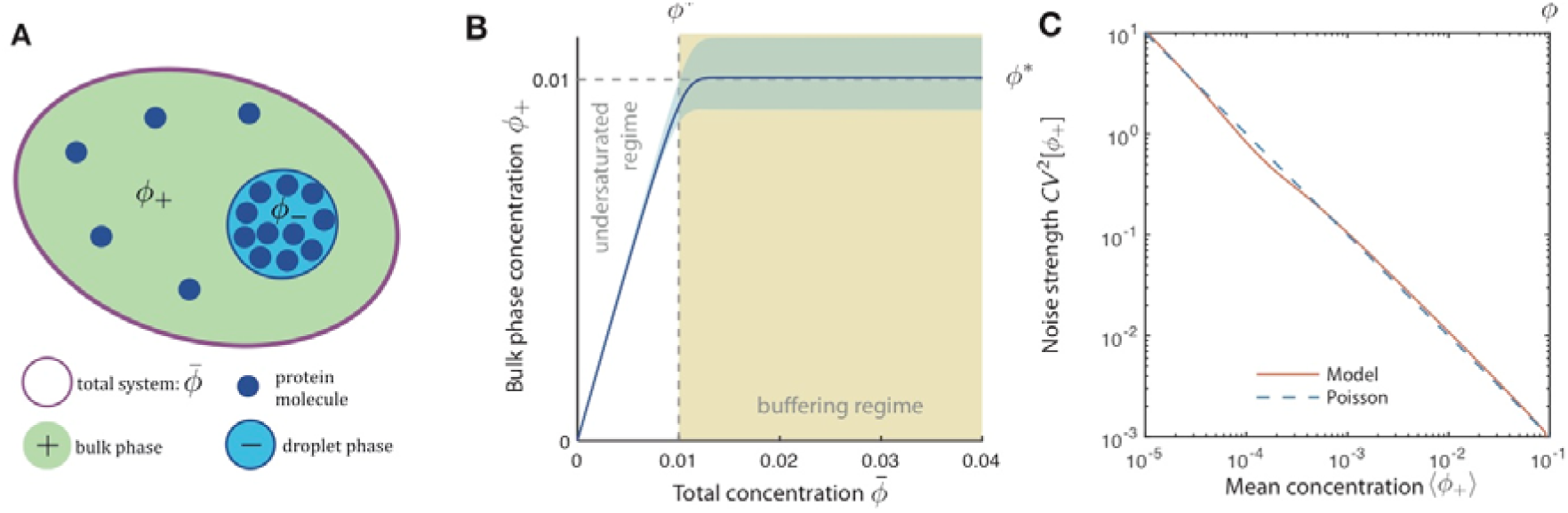
Concentration fluctuations in a phase-separating system at equilibrium. **(A)** Model schematic. We consider a binary mixture, consisting of a solvent and a protein (blue circles), which phase separates into a bulk (green) and droplet phase (cyan). We describe protein amounts in terms of normalized concentrations (volume fractions) defined by *ϕ* = *vc* with c as the concentration and v as the molecular volume of the protein. We denote the normalized concentrations of the bulk and droplet phase as well as of the total system as *ϕ*_+_, *ϕ*_-_and 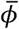, respectively. The system can be characterized in terms of its free energy, for which we considered contributions from both phases as well as the surface tension of the droplet (Supplementary text S.1.1.2). **(B)** Fluctuations in the bulk phase concentration *ϕ*_+_ as a function of the total protein concentration 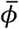. When 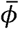 is greater than the threshold *ϕ*^*^, the system tends to phase separate, thus keeping *ϕ*_+_ close to *ϕ*^*^ with thermal fluctuations around it (blue shaded area indicate a range of 1 standard deviation around the mean). **(C)** Relationship between mean and noise strength of the bulk phase concentration *ϕ*_+_. The noise strength CV[*ϕ*_+_]^2^ was calculated from the considered equilibrium model for a fixed total concentration 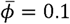. As soon as 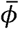 exceeds *ϕ*^*^ (buffering regime) the noise strength in the bulk phase concentration is approximately equal to the inverse of the mean, which resembles Poissonian noise (blue dashed line). A complete list of the model parameters can be found in Supplementary text Section S.3.

We next investigated if and to what extent the bulk phase noise deviates from the Poisson limit when variability is introduced by gene expression. For this purpose we constructed a model of gene expression^15^ accounting for stochastic transcription and translation, as well as extrinsic noise^16^ (Fig. 3A). By combining this model of gene expression with the mesoscopic theory of phase separation, we quantified the relation between fluctuations in the total protein concentration - characterized by its average 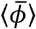 and noise strength 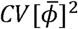 - and the corresponding fluctuations in the bulk phase protein concentration -characterized by (*ϕ*_+_) and CV[*ϕ*_+_]^2^. As a first step, we considered the limiting case where protein expression is slow with respect to the time scale of the droplet dynamics. In this limit, the bulk phase concentration is at quasi-equilibrium such that both (*ϕ*_+_) and CV[*ϕ*_+_]^2^ can be readily obtained from the equilibrium distribution of *ϕ*_+_ and the steady state distribution of 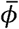 (Fig. 3B and Supplementary text S.1.2.2). Our analysis shows that as soon as the average total concentration 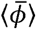 approaches the threshold concentration *ϕ*^*^, variations in the bulk phase start to decline due to the formation of droplets (Fig 3C). For larger 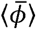, this effect becomes more and more pronounced until CV[*ϕ*_+_]^2^ approaches the Poisson limit (Fig. 3D). Furthermore, we found that gene expression noise can be suppressed almost entirely in this regime, while the remaining Poisson-like noise is due to thermal fluctuations of the droplets. We call this “buffering noise” (Supplementary text S.1.1.4). Therefore, our analysis predicts that gene expression noise can be suppressed by phase separation to the Poisson limit.

**Figure 3.**
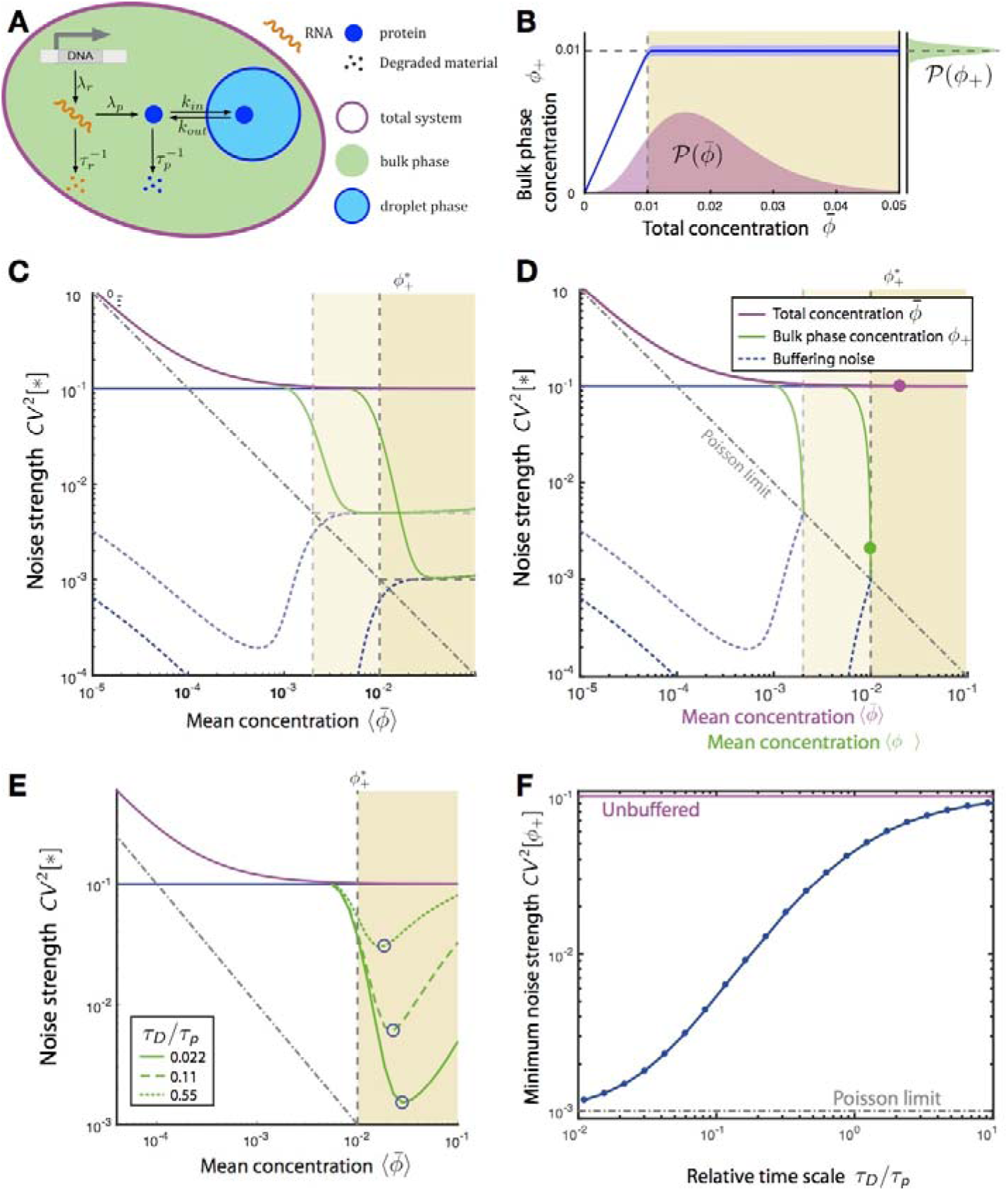
Buffering of protein expression noise by phase separation. **(A)** Model scheme. Fluctuations in protein concentration are described by a two-stage gene expression model accounting for stochastic production and degradation of RNA (λ_r_, r_r_) and protein (,λ_p_, r_p_). We consider protein degradation to take place in both phases. Partitioning of proteins into the bulk- and droplet phase is captured by stochastic exchange reactions with rates k_in_ and k_out_ derived from the thermodynamic model of phase separation. **(B)** Example distributions of the total and bulk phase concentrations 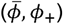. **(C,D)** Relationship between average concentration and noise strength for total (violet line) and bulk phase concentrations (green lines) for two different threshold concentrations 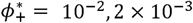, grey dashed lines) in the limit of slow protein dynamics. (C) Noise strength of the total and bulk phase concentration as a function of the mean total concentration. (D) Noise strength of the total and bulk phase concentration as a function of the mean total and mean bulk phase concentration, respectively. The green and violet circles indicate the noise strength corresponding to the example shown in (B). Blue dashed lines represent the amount of noise that is due to phase separation (Supplementary text S.1.1.4). **(E)** Dependency of noise buffering on time scales. The noise strength of the bulk phase is shown for three different relative time scales between protein expression and phase separation defined as the ratio between the average protein diffusion time and half live (r_D_/r_p_ = 0.022,0.11, .0.55). The blue circles indicate the minimum of the noise strength for the respective time scale. See also Fig. S4. **(F)** The minimal noise strength as indicated by circles in (E) for different time scale ratios r_D_/r_p_. The grey dashed lines indicate the noise strength that would be achieved in the absence of phase separation as well as the Poisson limit. For all subfigures we set 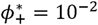 and CV[Λ_r_]^2^ = 0.1. A complete list of the model parameters can be found in Supplementary text S.3.

To study how the ability to buffer noise changes as the time scales of protein expression and phase separation become comparable, we developed a non-equilibrium model of phase separation that takes into account the different time scales associated with protein expression and phase separation. Fluctuations in the total- and bulk phase concentrations are then described by a master equation, from which we computed (*ϕ*_+_), CV[*ϕ*_+_]^2^, 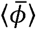 and 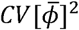 at steady state using van Kampen’s expansion^17^ (Supplementary text S.1.2.3). As before, our model exhibits a strong reduction of noise as soon as the system phase separates. Surprisingly, and in contrast to the quasi-equilibrium case, the bulk phase noise strength increases again as the mean total concentration increases (Fig 3E). This is because for large mean total concentrations, protein production is fast in comparison to the exchange of protein between the bulk and droplet phase. In this case, phase separation is too slow to effectively buffer rapid fluctuations in 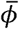.

In Fig. 3F, we plotted the minimal noise strength as a function of the time scale ratio of protein expression and phase separation. Here, the minimal noise strength was taken over all mean total concentrations as indicated by the blue circles in Fig. 3E for three different time scale ratios. This shows that the efficiency of noise buffering generally decreases with the inverse time scale of protein expression. However, a substantial reduction of noise can still be achieved if the time scales of phase separation and of protein expression are similar to each other.

We experimentally tested the prediction that phase separation can buffer protein expression noise by expressing the phase separating protein 2NT-DDX4^YFP^ inside HeLa cells, and examining protein concentration and spatial distribution using live-cell microscopy. The use of transient transfection allowed us to generate a broad range of protein expression levels across the cell population due to large variability in plasmid transfection efficiency (Fig. 4A). Similar to the previously described^14^ DDX4^YFP^, the 2NT-DDX4^YFP^ variant formed heterologous compartments inside nuclei of transfected HeLa cells (Fig. 4A). Duplicating the N-terminus reduced the protein’s threshold concentration and resulted in an increase in droplet-positive cells after transfection (Fig. S1). The droplets formed by 2NT-DDX4^YFP^ in the nuclei of transfected cells fused readily with each other, showed high internal recovery after photobleaching and exhibited hallmarks of Ostwald’s ripening over short time scales, which confirmed their liquid-like property (Fig. S2).

**Figure 4.**
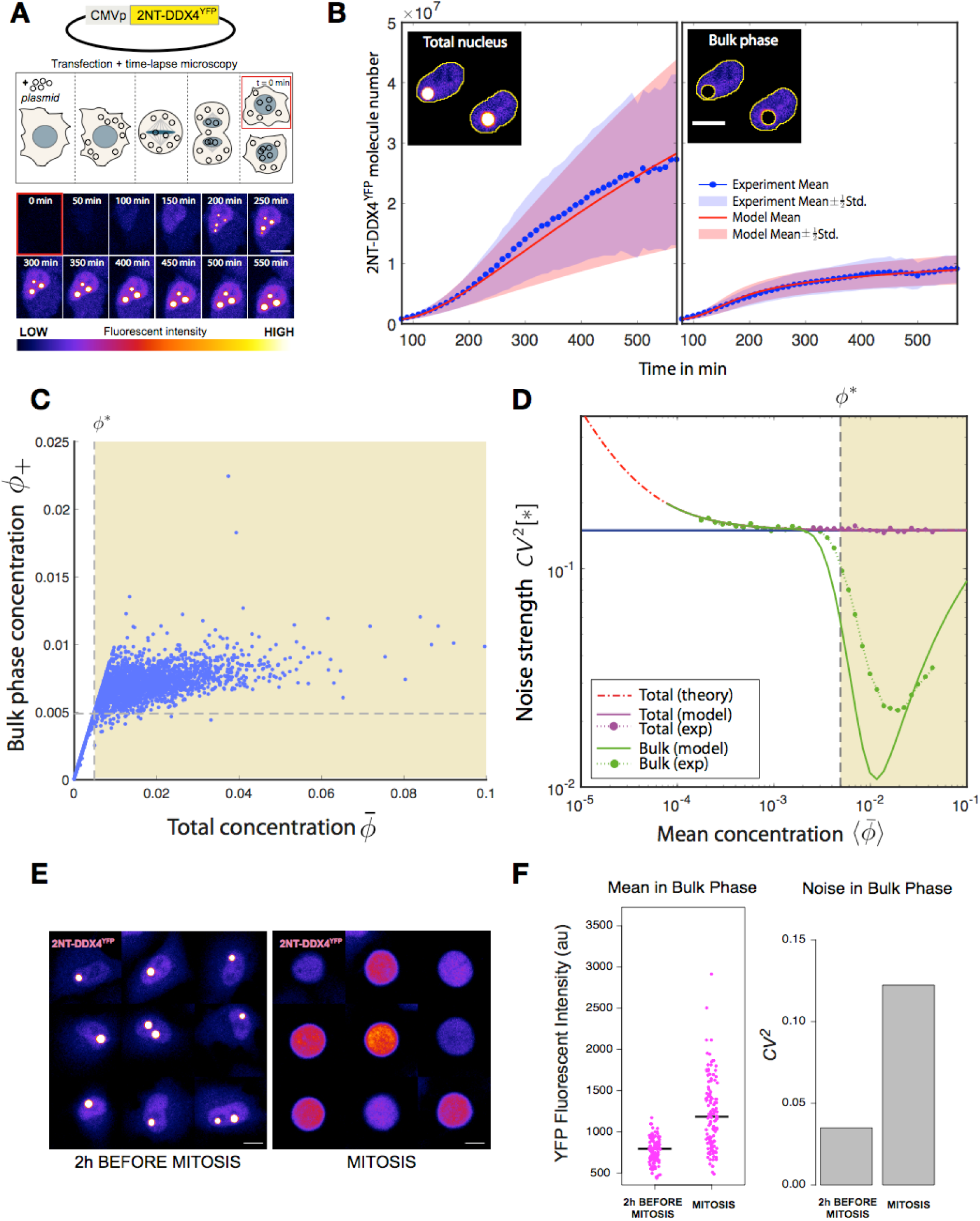
Efficient buffering of 2NT-DDX4^YFP^ expression noise in HeLa cells through phase separation. **(A)** Time-lapse microscopy of a typical cell initiating expression of 2NT-DDX4^YFP^ following transfection and plasmid incorporation. **(B)** Experimental time-lapse data and calibrated model. Kinetic parameters of 2NT-DDX4^YFP^ expression and phase separation were estimated using a moment-based inference approach (Supplementary text S.2.3). Solid lines represent average total and bulk phase protein number and shaded areas correspond to half a standard deviation. **(C)** Total versus bulk phase 2NT-DDX4^YFP^ concentrations quantified in a large number of cells 24hrs post transfection expressed as volume fractions. The dashed horizontal and vertical lines indicate the threshold concentration 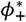 taken from the time-lapse measurements in (B). **(D)** Dependency between mean and noise strength of total and bulk phase 2NT-DDX4^YFP^ concentration. From the total population of cells, we randomly generated multiple subpopulations with different mean and noise strength of the total 2NT-DDX4^YFP^ concentration (main text and Supplementary text S.2.3). For each subpopulation, the noise strength in the bulk phase was predicted by the calibrated model and compared to the experimentally determined one. See also Fig. S5 **(E)** Montage of nine HeLa cells expressing 2NT-DDX4^YFP^ imaged before and during mitosis. **(F)** Quantification of mean 2NT-DDX4^YFP^ fluorescence intensity and noise in the bulk nucleoplasmic phase in 130 cells before and during mitosis. au = arbitrary units. *CV^2^* = squared coefficient of variation. Scale bar = 10 µm. Images were set to high contrast to demonstrate the change of intensity in the bulk phase.

To quantify the kinetics of protein concentration and phase separation, we used time-lapse fluorescence microscopy and measured the change of protein expression and droplet formation after transfection (Fig. 4A,B). Expression starts after the next mitosis, because the plasmid only incorporates into the nucleus after nuclear envelope reformation. We therefore identified post-mitotic cells in time-lapse videos and quantified total and bulk phase 2NT- DDX4^YFP^ concentrations in the nucleus (Fig 4A,B, see Materials and Methods). Over the first 200 minutes the protein concentration slowly increased and exhibited a single dilute phase. After 200 minutes, the protein began to form intra-nuclear droplets, which increased in number and size (Fig. 4A). This data allowed us to quantify the kinetic parameters of 2NT- DDX4^YFP^ expression and phase separation (see Materials and Methods and Fig. 4B and Supplementary text S.2.2).

We next determined the relationship between the mean concentration and the noise strength of 2NT-DDX4^YP^ in the bulk phase (in other words, the concentration outside the drops) and compared it to theory. For this purpose, we measured total and bulk phase 2NT-DDX4^YFP^ concentrations in a larger number of cells 24 hours after transfection (Fig. 4C). From the total pool of cells, we randomly selected subpopulations of cells which mimicked the statistics of the total protein concentration generated by our non-equilibrium model of protein expression. In this way, we obtained a series of subpopulations with different mean total concentrations. For each mean total-concentration we could then measure the mean and noise strength of bulk phase 2NT-DDX4^YFP^ protein concentration in the corresponding subpopulation (Supplementary text Section S.2.3). The resulting relationship between mean and noise strength in the bulk phase is in line with theory and confirms that gene expression noise is effectively buffered by phase separation in living cells (Fig. 4D). We note that the model underestimates the noise strength in the bulk phase. This is likely due to unknown additional sources of variability that are not captured by our simple model, such as cell-to-cell differences in the thermodynamic parameters or technical noise in image acquisition and processing. Our analysis shows that noise buffering is most effective for this particular protein when the mean total concentration is around 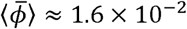 (corresponding to about 18 µM). At this point, the noise strength is reduced by almost seven-fold. Strikingly, for higher protein concentrations, the noise strength begins to increase again, as the model predicted.

We further confirmed the observation that reduction of noise is due to phase separation, by taking advantage of a previous observation that membraneless organelles dissolve during mitosis^18^. We followed over 100 individual droplet-containing cells through mitosis (Fig. S3) and quantified the fluorescence intensity in the nucleoplasm before and during mitosis. 2NT- DDX4^YFP^ droplets dissolve at the onset of mitosis (Fig. 4E) and this dissolution is associated with a three-fold increase of noise in the bulk phase when quantified across all the cells (Fig. 4F). This provides support for the idea that the reduction of noise is indeed due in part to phase separation.

System-wide single-cell studies^4,19^ have shown that the precision of gene regulation is constrained by biochemical noise. The magnitude of protein fluctuations generally decreases with average concentration, but even for highly expressed proteins, it rarely falls below a relative variation of around 10-20% (CV>0.1-0.2) depending on the organism and experimental condition^20,21^. How cells reliably process information and make decisions despite this limited precision has been a long-standing question. Previous studies have shown that transcriptional and post-transcriptional feedback mechanisms exhibit noise suppression^5,6^. However, the efficacy of these mechanisms is limited by the fact that they rely on cascades of biochemical steps, each of which can be slow and noisy itself. Phase separation, which happens post-translationally, is driven by thermodynamic principles that enable rapid and precise control of protein concentration, even in the presence of substantial noise. The data in this paper show that if the time scales of phase separation and protein expression are sufficiently separated, the bulk phase noise strength approaches the Poisson limit, irrespective of the noise strength in total protein concentration. Theory shows that noise buffering is generally less efficient when the dynamics of protein expression and phase separation become too similar. In cells, however, we expect the droplet dynamics to be fast in comparison to the production and turnover of protein, which typically takes place on the time scales of several hours. In the 2NT-DDX4^YFP^ system reported here, the time scales differed by about five-fold, which was sufficient to achieve significant noise buffering.

The present study provides a proof-of-principle using both theory and experiment that noise can be buffered by phase separation. Our experimental work uses an engineered system, which allows thorough quantification of concentration over a wide range of conditions, and comparison to theory. However, the same physical concepts should apply to endogenous phase separated compartments. Many such compartments have been identified recently but their biological function is often unclear^13,22,23^, and it seems likely that some of them serve as noise buffers that maintain protein concentrations at precise levels. As an example, paraspeckles are a ubiquitous compartment with no known function, and it has been shown that the bulk phase concentration of some paraspeckle proteins declines as paraspeckles form, suggesting a potential buffering role^23^.

Our study joins others that have proposed that spatial compartmentalization could be a mechanism to buffer noise^8^. For instance, in mammalian cells, delayed nuclear export of transcripts leads to reduced variability in cytoplasmic RNA^3^, as well as in yeast, where local clustering of protein can enhance the robustness of subcellular gradients^9^. In this paper we have demonstrated noise buffering for scaffold proteins, which themselves drive the formation of liquid compartments. However, many more proteins and RNAs partition into liquid compartments, for which similar ideas are likely to apply. These have been termed clients. Since clients are more abundant than scaffolds, this would expand the scope of the proposed mechanism to a substantial fraction of the genome. Indeed, about 30% of nuclear proteins are in condensates^24^.

In this paper, we have focused on the bulk phase, but because buffering is mediated by a modulation of droplet number and size, the protein concentration inside the droplets should be buffered as well. This could allow cells to create local microenvironments to enhance the fidelity of the reactions occurring inside of them. More systematic studies will be needed to understand the generality of noise buffering by phase separation in endogenous biological systems.

## Materials and Methods

### Protein purification

The recombinant DDX4^YFP^ and 2NT-DDX4^YFP^ proteins were expressed from an inducible bacterial expression vector where it was sub-cloned with an N-terminal MBP tag and C-terminal 6xHis tag, both flanked by a 3C (‘PreScission’) site. *E. coli* BL21 cells carrying a pRARE plasmid were transfected with the expression plasmid and grown overnight under antibiotic selection (Kanamycin + Chloramphenicol) in LB medium. The next day, bacteria were grown to OD=0.6 in TB medium, then moved from 37 to 20 °C for 1 h and induced overnight with 200 µM IPTG. The next day cells were harvested by centrifugation, resuspended in lysis buffer (50 mM Tris-HCl pH = 8.0, 500 mM NaCl, 1 mM DTT, 0.5 µg/ml benzonase, 10 µg/ml lysozyme, 1mM MgCl_2_, 1:1000 Protease Inhibitor Coctail (bimake.com)) and further disrupted by sonication, followed by centrifugation for 45 min at 4 °C. The clarified lysate was loaded on gravity column previously packed with Ni-NTA agarose beads (QIAGEN) and equilibrated with working buffer (50 mM Tris-HCl pH = 8.0, 500 mM NaCl, 1 mM DTT). After loading, the columns were washed with 3 column volumes of working buffer and eluted with Ni-elution buffer (working buffer + 250 mM Imidazole). The sample was then passed through Amylose Resin (NEB) with a similar protocol and eluted with MBP-elution buffer (working buffer + 10mM maltose). The terminal MBP and 6xHis tags were cleaved by incubation with 3C PreScission protease at a 1:100 ratio in room temperature overnight. The protein was purified with gel filtration chromatography (Superdex-200; GE Healthcare) equilibrated with storage buffer (20 mM Tris-HCl pH = 8.0, 500 mM NaCl, 1 mM DTT). The fractions corresponding to the expected protein size and fluorescent emission were pooled and concentrated to 50 – 200 µM protein, flash-frozen in liquid nitrogen and stored at −80°C.

### *In vitro* assay

The dilutions were made first in high salt buffer (500 mM NaCl, 20 mM Tris-HCl pH = 8.0, 1 mM DTT) and the concentration of protein and NaCl were dropped concomitantly by diluting with no-salt buffer (20 mM Tris-HCl pH = 8.0, 1 mM DTT) to achieve the final protein concentration used in the droplet assays. Samples were imaged in self-made microchambers where 1 µl drop of sample was pressed between glass and a cover-slide separated and sealed by a spacing made by double-sided scotch tape. Imaging was done using Andor spinning disk microscope equipped with 488nm solid-state laser, 40x Silicon Objective and iXon EM+ DU-897 BV back illuminated EMCCD camera. For the centrifugation assay, the samples were spun at 15.000 rpm for 10 min and concentration in the supernatant determined by measuring absorbance at 280 nm using a NanoDrop ND-1000 spectrophotometer (Thermo Scientific).

### Cell culture and transfection

All constructs used for transfection were derived from a plasmid expressing recombinant DDX4^YFP^ protein that was a gift from Timothy Nott. In 2NT-DDX4^YFP^ construct the entire N-terminal region encoding 247 amino acids was inserted in-frame upstream of the start codon effectively generating a recombinant product with a duplicated N-terminus. The cell line used was a HeLa Kyoto line cultured in high glucose DMEM (Gibco, catalogue # 11995-065) containing 0.5 mg/ml of Penicillin-Streptomycin (Gibco, catalogue # 10378-016) and 10 % fetal bovine serum (Gibco, catalogue # 26140-079). For the time-lapse microscopy cell line constitutively expressing H2B:mCherry from a plasmid kept at stable level by addition of Blasticidin to the culture media was used (Life Technologies, 2 μg/ml). Cells were seeded in 8-well µ-slide (Ibidi, catalogue # 80826) 24 hours prior to transfection. Transfection was done using plasmid purified with PureLink HiPure Plasmid Filter Midiprep Kit (Invitrogen, catalogue # K2100-15) with 200 ng of DNA/well and Lipofectamine 2000 Transfection Reagent (Invitrogen, catalogue # 11668-019) following manufacturers instruction. Opti-MEM (Gibco, catalogue # 11058-021) supplemented with 10 % fetal bovine serum (Gibco, catalogue # 26140-079) was used for cell culture during and following the transfection.

### Microscopy

Long term time-lapse imaging was started four hours after transfection using Olympus IX83 spinning disk confocal microscope equipped, with Andor iXon 888 Ultra camera, temperature control chamber set at 37 °C and 5 % CO_2_ and using 40x silicon immersion objective. A complete Z-stack at each position in YFP and mCherry channel was acquired every 10 minutes for a total of >11 hours. Same hardware and setting were used for collecting images at the steady state at 24 – 28 hours post transfection. Hoechst dye was added directly prior to imaging at 200 ng/ml final concentration. A complete Z-stack at >300 positions was acquired in YFP and Hoechst channels.

### Image analysis

Images were analyzed using a custom pipeline made with Fiji^25^. Briefly, the DNA signal emanating from the nucleus in the blue channel was used to threshold the nuclei and the droplets were segmented based on their fluorescence using a fixed minimal threshold for all images. The mean intensity of the segmented nucleus and the bulk nucleoplasm (nucleus minus droplets) were converted to concentrations using a calibration curve made using serial dilutions of purified 2NT-DDX4^YFP^ protein imaged during the same imaging session. Concentration of the purified protein was estimated using absorbance at 280 nm and a linear fit was made to establish the relationship of the fluoresce intensity measure at the microscope to the protein concentration.

