## Supplementary text for "Phase separation provides a mechanism to reduce noise in cells"

**3 Parameter values used for the figures in the main paper 14**

### 1 Theory of concentration fluctuations in mesoscopic droplet systems

In this section, we will develop a theoretical model to study the influence of liquid-liquid phase separation on gene expression variability. In Section 1.1, we will first construct a simple mesoscopic model of phase separation based on the equilibrium physics of binary mixtures. Subsequently, we will extend the model to account for variability in the composition of this mixture due to gene expression noise. In Section 2 we show how the model was used to analyze and interpret our experimental single-cell data.

#### 1.1 Minimal model of phase separation in a binary mixture

We consider a system of a fixed volume $V_{tot}$ (e.g. the volume of the cell or the nucleus) which is filled by a binary mixture. Each of the two components of the mixture can represent a single molecule or a bundle of different molecules in a fixed ratio that form a unit. For the sake of clarity, we refer to one component as 'solvent' and to the other as 'protein' in the following. The two components can phase separate by forming a droplet of volume $V_-$ which is surrounded by a bulk phase of volume $V_+ = V_{tot} - V_-$ (Fig. ST1). Thereby the total number of proteins $\bar{N}$ is split up between the two phases such that the phases differ by their concentrations ($c_\pm = N_\pm/V_\pm$) of the protein. Here $N_-$ denotes the number of molecules or units of the component that is enriched in the droplet, i.e. the protein, inside the droplet, while $N_+$ is the number of molecules in the surrounding bulk phase. With the effective molecular volume, we can define a volume fraction occupied by the protein (component) as $\phi = vN/V$. As explained in the main text volume fractions can, therefore, be understood as normalized concentrations, with $\phi = 0$ equal to a system without any protein molecule and $\phi = 1$ for a system completely filled with the protein component. We will focus on the comparison of noise in the volume fraction of the whole system, $\bar{\phi} = v\bar{N}/V_{tot}$, to the noise of volume fraction of the bulk phase, $\phi_+ = vN_+/V_+$. We aim to understand if and to which extent such a system is able to keep the volume fraction of the bulk phase close to a fixed value. This ability is what we refer to as *buffering*.

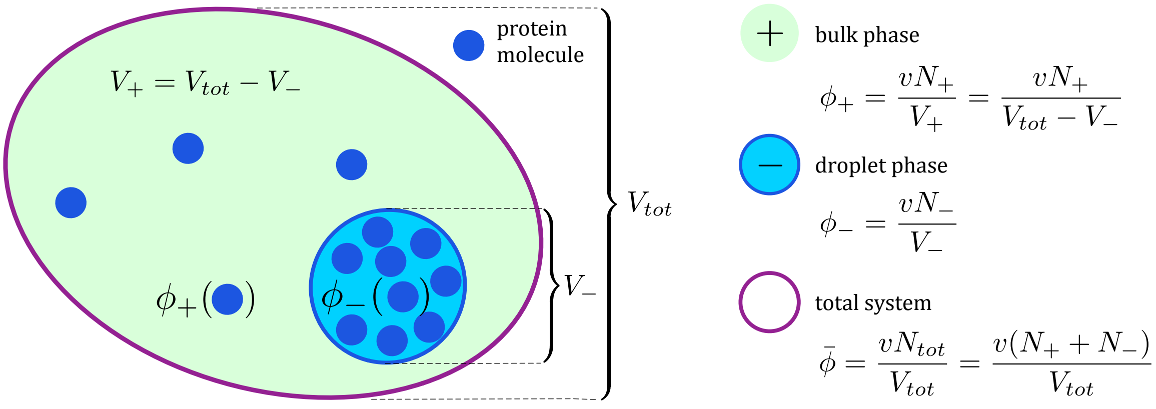

**Figure ST1:** Minimal model of phase separation in a binary mixture.

##### 1.1.1 Free energy of a binary mixture

To describe the thermodynamic properties of such a mesoscopic system containing a single droplet we have to consider its free energy. If the respective volume fractions within the droplet and in the solvent are homogeneous, the free energy $F$ of the system is given by

$$F = V_- f_-(\phi_-) + (V_{tot} - V_-) f_+(\phi_+) + \gamma A \tag{1}$$

where $f_\pm(\phi_\pm)$ are the free energy densities of the droplet phase and the bulk phase respectively and $\gamma$ and $A$ are the surface tension and the area of the droplet-bulk interface. We consider the case of a droplet that is entirely filled with one component

$$\phi_- = vN_-/V_- = vc_- = 1\ , \tag{2}$$

such that we can write the free energy density of the droplet phase as

$$f_{-}(\phi_{-}) = -\frac{\mu}{v} \ , \tag{3}$$

with $\mu$ as a relative chemical potential. From eq. (2) also follows $v = 1/c_{-}$. With eq. (2) and $\bar{N} = N_{+} + N_{-}$ we can express the volumes of the two phases as:

$$V_{-}\left(N_{+}, \bar{N}\right) = v\left(\bar{N} - N_{+}\right)$$
$$V_{+}\left(N_{+}, \bar{N}\right) = V_{tot} - v\left(\bar{N} - N_{+}\right)$$

Furthermore, we consider the droplet to be spherical such that

$$A = 4\pi\left(\frac{3}{4\pi}vN_{-}\right)^{\frac{2}{3}} \ . \tag{4}$$

The free energy density of the bulk phase is considered to contribute as an ideal gas, i.e.

$$f_{+}\left(\phi_{+}\right) = \frac{k_B T}{v}\left[\phi_{+}\ln\left(\phi_{+}\right) - \phi_{+}\right] \ , \tag{5}$$

where $k_B$ denotes the Boltzmann constant and $T$ the temperature. The free energy of the whole system can then be written as

$$F\left(N_{+}, \bar{N}\right) = -\mu\left(\bar{N} - N_{+}\right) + k_B T N_{+}\left(\ln\left(\frac{N_{+}}{V_{tot}/v - \bar{N} + N_{+}}\right) - 1\right) + \gamma\left(36\pi\right)^{\frac{1}{3}}\left(v\left(\bar{N} - N_{+}\right)\right)^{\frac{2}{3}} . \tag{6}$$

Equivalently, the free energy can be expressed in terms of volume fractions

$$\hat{F}\left(\phi_{+}, \bar{\phi}\right) = -\mu\frac{V_{tot}}{v}\frac{\left(\bar{\phi} - \phi_{+}\right)}{\left(1 - \phi_{+}\right)} + k_B T\frac{V_{tot}}{v}\frac{(1 - \bar{\phi})\phi_{+}}{\left(1 - \phi_{+}\right)}\left(\ln\left(\phi_{+}\right) - 1\right) + \gamma\left(36\pi\right)^{\frac{1}{3}}\left(V_{tot}\frac{\left(\bar{\phi} - \phi_{+}\right)}{\left(1 - \phi_{+}\right)}\right)^{\frac{2}{3}} . \tag{7}$$

##### 1.1.2 Thermal fluctuations at equilibrium: Boltzmann statistics

From here on we follow the convention to use upper case letters for random variables and lower case letters for their realization. To analyze the influence of thermal fluctuations on the buffering properties of the system, we calculate the partition function:

$$Z_{\bar{n}} = \sum_{n_{+}=0}^{\bar{n}} \exp\left(-\frac{F\left(n_{+}, \bar{n}\right)}{k_B T}\right) \ , \tag{8}$$

such that the Probability $\mathcal{P}_{eq}$ to find the system in a state with $n_{+}$ molecules outside the droplet is given by the Boltzmann distribution

$$\mathcal{P}_{eq}\left(N_{+} = n_{+}\right) = \frac{1}{Z_{\bar{n}}}\exp\left(-\frac{F\left(n_{+}, \bar{n}\right)}{k_B T}\right) \ . \tag{9}$$

Equivalently, this distribution can be expressed in terms of volume fractions, i.e., $\mathcal{P}_{eq}\left(\phi_{+}\right) = \mathcal{P}_{eq}\left(N_{+} = \phi_{+}V/v\right)$. With eq. (9), we can determine the mean and variance of the volume fraction of the bulk phase as:

$$\langle\Phi_{+}\rangle = \sum_{n_{+}=0}^{\bar{n}} \mathcal{P}_{eq}\left(N_{+} = n_{+}\right)\phi_{+}\left(n_{+}, \bar{n}\right) \tag{10}$$

$$\sigma^2\left(\Phi_{+}\right) = \left\langle\left(\Phi_{+} - \langle\Phi_{+}\rangle\right)^2\right\rangle = \sum_{n_{+}=0}^{\bar{n}} \mathcal{P}_{eq}\left(N_{+} = n_{+}\right)\left[\phi_{+}\left(n_{+}, \bar{n}\right) - \langle\Phi_{+}\rangle\right]^2 . \tag{11}$$

Throughout this text, we denote means and variances of random variables as $\langle\,\cdot\,\rangle$ and $\sigma^2\left(\,\cdot\,\right)$, respectively. Means and variances of bulk phase are illustrated in Fig. ST2 for different $\bar{\phi}$.

For the sake of readability, we drop the explicit dependence of the volume fraction on the protein numbers in the following. However, the reader should keep in mind that with discrete protein numbers, also the corresponding volume fractions are discrete-valued.

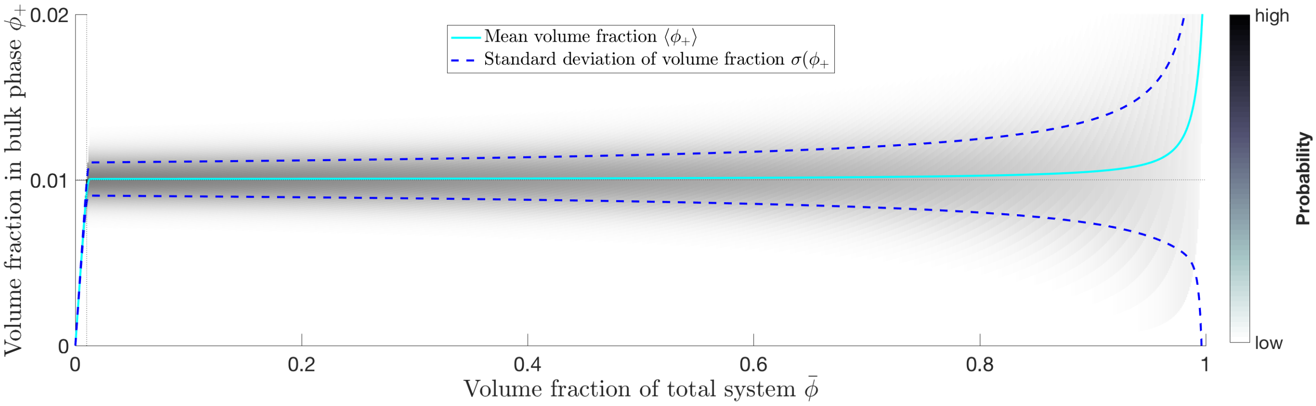

**Figure ST2:** Thermal fluctuations at equilibrium. The grey shaded areas illustrate the probability to find a certain volume fraction in the bulk phase. The solid cyan line indicates the mean and the dashed blue lines indicate a standard deviation around the mean, i.e. $\langle\Phi_+\rangle \pm \sigma(\Phi_+)$. The model parameters are set to the same values as for Figure 2 in the main text and are listed in Tab. 3 on page 14.

##### 1.1.3 Analytical characterization of thermal fluctuations

To find an analytical expression for the concentration fluctuations, we approximate the Boltzmann distribution using the saddle point method. To this end, we interpret the protein volume fractions as continuous quantities and expand the free energy around its minimum up to order two. The global minimum of the free energy, (if it exists, i.e. if $\bar{\phi}$ is sufficiently large and phase separation is favored) obeys

$$
\begin{aligned}
0 =& \frac{\partial \hat{F}\left(\phi_+, \bar{\phi}\right)}{\partial \phi_+} \\
=& \frac{V_{tot}(1-\bar{\phi})}{v(1-\phi_+)^2}\left(\mu + k_B T\left(\ln\phi_+ - \phi_+\right) + 2\gamma H\right) \ ,
\end{aligned}
\tag{12}
$$

where $H = \frac{\partial A}{\partial V_-}$ is the mean curvature. When the contribution of the interface energy becomes negligible (i.e. if $\gamma$ is small, or if the system approaches the thermodynamic limit), eq. (12) is satisfied if:

$$
0 = \mu + k_B T\left(\ln\phi_+ - \phi_+\right). \tag{13}
$$

This provides and implicit relation for the saddle point of the Boltzmann distribution and we refer to its solution as $\phi^*$. The latter can be expressed in terms of Lambert W function:

$$
\phi^* = -W\left(-e^{\frac{-\mu}{k_B T}}\right) \ . \tag{14}
$$

Since this function is invertible on the domain $\phi^* \in [0,1]$, we can replace the dependency of the Boltzmann distribution on $\mu$ by $\phi^*$. The new parameter $\phi^*$ has an important physical interpretation. In particular, it corresponds to the volume fraction above which a mesoscopic system is expected to exhibit two distinct phases. In the thermodynamic limit, the volume fractions in the bulk and droplet phase change discontinuously at $\phi^*$ owing to a thermodynamic phase transition. In the following, we will refer to $\phi^*$ as *threshold concentration.*

In order to characterize the fluctuations in the bulk phase in the phase separating regime, we further calculate the second derivative of the free energy at the saddle point $\phi^*$,

$$
\left(\frac{\partial}{\partial\phi_+}\right)^2 \hat{F}\left(\phi_+, \bar{\phi}\right)\Bigg|_{\phi_+=\phi^*} = k_B T \frac{V_{tot}\left(1-\bar{\phi}\right)}{v\phi^*\left(1-\phi^*\right)} \ , \tag{15}
$$

where we again neglected contributions from the surface tension. We can now approximate the Boltzmann distribution with the saddle point method as a Gaussian

$$
\widetilde{p}_{eq}\left(\phi_+\right) = \frac{1}{\widetilde{Z}_{\bar{\phi}}} \exp\left(-\frac{V_{tot}\left(1-\bar{\phi}\right)}{2v\phi^*\left(1-\phi^*\right)}\left(\phi_+ - \phi^*\right)^2\right) \ , \tag{16}
$$

where $\widetilde{p}_{eq}$ is now a probability density and $\widetilde{Z}_{\bar{\phi}}$ is the corresponding normalization constant. The mean and variance of (16) are given by

$$\langle \Phi_+ \rangle = \phi^* \qquad \text{and} \qquad \sigma^2(\Phi_+) = \frac{v\phi^*(1-\phi^*)}{V_{tot}\left(1-\bar{\phi}\right)} \tag{17}$$

and the squared coefficient of variation becomes

$$\eta^2(\Phi_+) = \frac{\sigma^2(\Phi_+)}{\langle \Phi_+ \rangle^2} = \frac{v(1-\phi^*)}{V_{tot}\left(1-\bar{\phi}\right)\phi^*} \,. \tag{18}$$

Throughout this text, we denote the coefficient of variation by $\eta(\,\cdot\,) = \sigma(\,\cdot\,)/\langle\,\cdot\,\rangle$ and define by $\eta^2(\,\cdot\,)$ the *noise strength* of the system, a commonly used metric to assess concentration variability.

When both the threshold concentration and total volume fraction are small $\left(\phi^* < \bar{\phi} \ll 1\right)$, the above expression simplifies to

$$\eta^2(\Phi_+) \approx \frac{v}{V_{tot}\phi^*} \,. \tag{19}$$

Since $n_- \leq \bar{n}$ we can without further assumptions approximate $\phi^*$ by expanding $\phi_+$ in $n_-$ as

$$\begin{aligned} \phi^* = \langle \Phi_+ \rangle &= \left\langle \frac{N_+}{V_{tot}/v - N_-} \right\rangle \\ &= \left\langle \frac{N_+}{V_{tot}/v} + \frac{N_+N_-}{(V_{tot}/v)^2} + \mathcal{O}\left(N_-^2\right) \right\rangle \\ &= \frac{\langle N_+ \rangle}{V_{tot}/v} + \frac{\langle N_+N_- \rangle}{(V_{tot}/v)^2} + \left\langle \mathcal{O}\left(N_-^2\right) \right\rangle \,. \end{aligned} \tag{20}$$

With eqs. (19) and (20), the noise strength is approximately given by

$$\eta^2(\Phi_+) \approx \frac{1}{\langle N_+ \rangle} \,. \tag{21}$$

This resembles the mean-noise relationship of a Poisson-distributed random variable.

###### 1.1.4 Ensembles of phase separating systems with different molecule numbers

So far, we have studied fluctuations of phase separating systems for which the total protein number was considered to be fixed. We will next extend our analysis to the case where the total protein concentration is a random variable itself. This is an important intermediate step that will later allow us to study how fluctuations caused by gene expression are affected by phase separation (Section 1.2).

To address this scenario, we consider an ensemble of independent phase separating systems at equilibrium, each having identical material parameters, but different total protein amounts. To capture the variability in the total protein amount across the ensemble, we describe the total protein volume fraction as a random variable $\bar{\Phi} \sim \mathcal{P}\left(\bar{\phi}\right)$, with $\mathcal{P}\left(\bar{\phi}\right)$ as an arbitrary, discrete probability distribution. Our goal is now to relate the variability in the total protein amount (characterized by $\mathcal{P}\left(\bar{\phi}\right)$) to the variability in the bulk phase (characterized by a corresponding distribution $\mathcal{P}(\phi_+)$). If the ensemble is at equilibrium, the distribution $\mathcal{P}(\phi_+)$ can be readily obtained from the Boltzmann distribution in eq. (9). Since the total protein amount $\bar{\Phi}$ is now a random variable, the Boltzmann distribution associated with an individual phase separating system is now interpreted as a conditional probability distribution $\mathcal{P}_{eq}\left(\phi_+|\bar{\Phi} = \bar{\phi}\right)$, where $\bar{\phi}$ is the particular realization of $\bar{\Phi}$ corresponding to that system. In order to compute $\mathcal{P}(\phi_+)$ we have to average this conditional distribution over all possible realizations of $\bar{\Phi}$, i.e.,

$$\mathcal{P}(\phi_+) = \sum_{\bar{\phi}} \mathcal{P}_{eq}\left(\phi_+|\bar{\Phi} = \bar{\phi}\right)\mathcal{P}\left(\bar{\phi}\right) . \tag{22}$$

With this expression, we can now compare the noise strength of the total protein amount and the bulk phase by analyzing $\mathcal{P}\left(\bar{\phi}\right)$ and $\mathcal{P}(\phi_+)$, respectively.

Moreover, we can use the law of total variation to decompose the noise strength of the bulk phase into

$$\eta^2(\Phi_+) = \underbrace{\frac{\sigma^2\left(\langle \Phi_+|\bar{\Phi}\rangle\right)}{\langle \Phi_+ \rangle^2}}_{\text{ensemble noise}} + \underbrace{\frac{\left\langle \sigma^2\left(\Phi_+|\bar{\Phi}\right)\right\rangle}{\langle \Phi_+ \rangle^2}}_{\text{buffering noise}} . \tag{23}$$

If the systems in the ensemble are likely below the threshold concentration $\phi^*$, the bulk phase concentration $\phi_+$ is approximately equal to $\bar{\phi}$. In this case, $\sigma^2\left(\Phi_+|\bar{\Phi}=\bar{\phi}\right)$ and $\sigma^2\left(\langle\Phi_+|\bar{\Phi}\rangle\right)\approx\sigma^2\left(\bar{\Phi}\right)$, which means that the second term in (23) vanishes and that the first term explains all the variability in the bulk phase. In this scenario, the variability in the total concentration cannot be suppressed and is entirely inherited to the bulk phase. If, on the other hand, all systems in the ensemble would be above $\phi^*$, we know by eq. (17) that the expected bulk phase volume fraction is widely independent of the total volume fraction $\bar{\phi}$, such that the first term is close to zero and the second term captures the expected thermal fluctuations around the threshold concentration. In this sense, the first term of eq. (23) describes the variance due to ensemble variability, while the second term describes the noise introduced by phase separation itself. We thus refer to the respective contributions as *ensemble noise* and *buffering noise*.

##### 1.1.5 Stochastic dynamics of phase separating systems at equilibrium

In our discussion above, we have described concentration fluctuations in terms of probability distributions over the total protein and bulk phase concentration. We will next investigate the temporal dynamics of phase separating systems at equilibrium. To this end, we will derive rate functions of protein exchange between the bulk and droplet phase, such that the resulting fluctuations are consistent with the equilibrium fluctuations of mesoscopic droplet systems described in Section 1.1.2. These rate functions will be instrumental later in Section Section 1.2 to link our theory of phase separation to gene expression noise.

For a fixed total protein number $\bar{N}$, the dynamic partitioning of protein between the bulk and droplet phase can be described by stochastic exchange reactions

$$\mathrm{N}_+ \underset{k_{in}}{\overset{k_{out}}{\rightleftharpoons}} \mathrm{N}_-,$$

with $N_\pm$ as the bulk and droplet phase protein and $k_{in}$ and $k_{out}$ as the respective reaction rates. These rates determine how likely each reaction takes place per unit time and thus set the time scale of phase separation. Our goal is to determine explicit expressions for $k_{in}$ and $k_{out}$ such that they are consistent with our equilibrium model of phase separation. At equilibrium, the rates satisfy the detailed balance condition:

$$k_{out}\left(n_+,n_-\right)\mathcal{P}_{eq}\left(n_+\mid\bar{N}=n_++n_-\right)=k_{in}\left(n_++1,n_--1\right)\mathcal{P}_{eq}\left(n_++1\mid\bar{N}=n_++n_-\right) \tag{24}$$

or in terms of the free energy:

$$k_{out}\left(n_+,n_-\right)=k_{in}\left(n_++1,n_--1\right)\exp\left(-\frac{\Delta F\left(n_+,n_-\right)}{k_BT}\right)\,, \tag{25}$$

with $\Delta F\left(n_+,n_-\right)=F\left(n_++1,n_++n_-\right)-F\left(n_+,n_++n_-\right)$. Therefore, if one of the two rates is given, the reverse rate can be determined by the Boltzmann factor. Here we consider the rate $k_{in}$ to be diffusion-limited such that

$$k_{in}\left(n_+,n_-\right)=\frac{n_+}{\tau_d\left(n_-\right)}=\frac{6D\,n_+}{\left(V_{tot}-vn_-\right)^{2/3}}=\frac{6DV_+^{1/3}}{v}\phi_+, \tag{26}$$

with $\tau_d$ as the average time it takes for a molecule to diffuse through a volume $V_+$ and $D$ as the diffusion constant. The rate from (26) also provides a simple way to characterize the time scale of phase separation. In particular, we can define it as the diffusion time of a single protein through the total system volume $V_{tot}$, i.e., $\tau_D=\tau_d(0)=V_{tot}^{2/3}/(6D)$. We will later make use of $\tau_D$ when we analyze how time scales affect noise buffering by liquid-liquid phase separation.

#### 1.2 Suppression of gene expression noise by liquid-liquid phase separation

Based on our theoretical considerations from the previous sections, we are now ready to investigate how phase separation can buffer gene expression variability. We will first introduce a canonical model of stochastic gene expression and subsequently show how this model can combined with our model of liquid-liquid phase separation.

##### 1.2.1 Two-stage model of gene expression with extrinsic noise

We describe gene expression by a two-stage model, where mRNA and protein copy numbers, denoted by $N_r$ and $N_p$, are described by coupled birth-and-death processes

$$\begin{aligned} \emptyset &\xrightarrow{\lambda_r} \mathrm{N}_r \xrightarrow{N_r/\tau_r} \emptyset \\ \emptyset &\xrightarrow{\lambda_p N_r} \mathrm{N}_p \xrightarrow{N_p/\tau_p} \emptyset \end{aligned} \tag{27}$$

with $\lambda_r$ and $\lambda_p$ as the rate constants associated with transcription and translation and $\tau_r$ and $\tau_p$ as mRNA and protein half-life times. The noise strength of $N_p$ at steady state is given by [1]

$$\eta^2\left(N_p\right) = \frac{1}{\langle N_p \rangle} + \frac{1}{\langle N_r \rangle}\frac{\tau_r}{\tau_r + \tau_p}. \tag{28}$$

Additional to intrinsic fluctuations of transcription and translation, gene expression is significantly affected by so-called extrinsic variability[2]. The latter stems from cell-to-cell differences in factors affecting gene expression such as ribosome copy numbers or the ATP availability. In the experimental system considered here, the dominant source of extrinsic variability stems from plasmid copy variations, which can be incorporated into the model by considering the transcription rate to be randomly distributed across cells, i.e., $\Lambda_r \sim p(\lambda_r)$. We want to remark that while our analysis focuses on this particular model of gene expression, it is straightforward to extend to other scenarios.

For the two-stage model with random transcription rate, one can show that the noise strength is given by

$$\begin{aligned} \eta^2\left(N_p\right) &= \eta^2\left(\langle N_p|\Lambda_r\rangle\right) + \left\langle \eta^2\left(N_p|\Lambda_r\right)\right\rangle \\ &= \eta^2\left(\Lambda_r\right) + \left(1 + \frac{\lambda_p}{\tau_p^{-1} + \tau_r^{-1}}\right)\frac{1}{\langle N_p\rangle}. \end{aligned} \tag{29}$$

Throughout this study, we consider the transcription rate $\lambda_R$ to be Gamma-distributed with shape- and rate parameters $\alpha$ and $\beta$, i.e.

$$\Lambda_r \sim \Gamma(\alpha,\beta) = \frac{\beta}{\Gamma(\alpha)}\left(\beta\lambda_r\right)^{\alpha-1} e^{-\beta\lambda_r}\ . \tag{30}$$

In this case, the mean and noise strength of the transcription rate is given by $\langle \Lambda_r \rangle = \alpha/\beta$ and $\eta^2\left(\Lambda_r\right) = \alpha^{-1}$, respectively. Thus, the amount of extrinsic noise in the system is set by $\alpha$, while the rate parameter $\beta$ determines the average protein expression level, i.e., $\langle N_p\rangle = \langle \Lambda_r\rangle \tau_r \lambda_p \tau_p$. This provides a way to calculate the mean-noise relationship of the gene expression network for a fixed extrinsic noise level by evaluating (29) for different $\beta$. The resulting relationship shows the characteristic inverse scaling of the noise strength with mean concentration. For high concentrations, the SCV saturates at a certain level due to extrinsic noise (Fig. ST3). This mean-noise relationship is a hallmark of stochastic gene expression [3, 4] and we will next discuss how it is affected by liquid-liquid phase separation.

##### 1.2.2 Limit of fast phase separation dynamics

We will now investigate how variations in protein concentrations can be suppressed by liquid-liquid phase separation. As a first step, we consider a simplified scenario that considers phase separation to be fast on the time scale of protein expression. More precisely, we consider phase separation to equilibrate instantaneously upon changes in the total protein concentration. This in turn allows us to apply the equilibrium theory presented in Section 1.1.4, whereas the ensemble distribution $\mathcal{P}\left(\bar{\phi}\right)$ now corresponds to the stationary distribution of the stochastic gene network in (eq. (27)). In general, this distribution can be computed by numerically solving a master equation but also analytical solutions exist under certain conditions. If the mRNA fluctuates much faster than the protein, for instance, the stationary distribution over protein copy numbers is given by a negative binomial distribution $\mathcal{P}\left(\bar{N} = \bar{n}\right) = \mathcal{NB}(r,p)$ with

$$\mathcal{P}(\bar{N} = \bar{n}) = \binom{\bar{n} + r - 1}{\bar{n}} p^r (1-p)^{\bar{n}} \tag{31}$$

with parameters

$$r = \frac{1}{\eta^2\left(\Lambda_r\right)} \qquad p = \frac{r}{r + \langle \bar{N}\rangle}\ . \tag{32}$$

The corresponding distribution over volume fractions is then given by $\mathcal{P}\left(\bar{\Phi} = \bar{\phi}\right) = \mathcal{P}\left(\bar{N} = \bar{\phi}V_{tot}/v\right)$. Figure ST3 illustrates the resulting mean-noise relationships for the considered model. More importantly, we show how this mean-noise relationship is affected by liquid-liquid phase separation in Figure ST4.

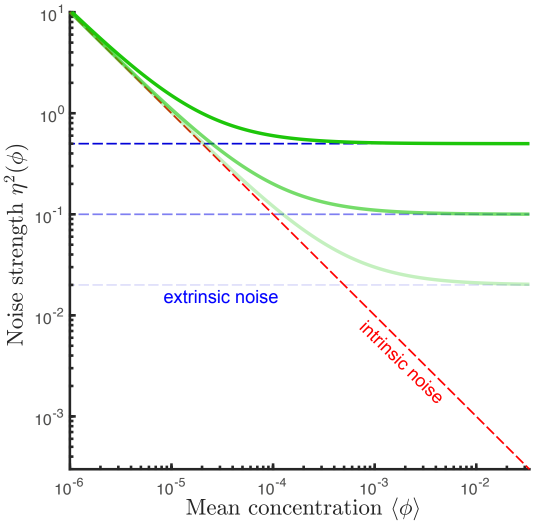

**Figure ST3:** Mean-noise relationship (green solid lines) in stochastic gene expression described by the negative binomial distribution for three different levels of extrinsic noise, $\eta^2(\Lambda_r) = [0.02,\ 0.1,\ 0.5]$, indicated by the blue dashed lines. The remaining parameters are set to the same values as for Figure 3B and 3C in the main paper and are noted in Tab. 4.

##### 1.2.3 Influence of fluctuating protein concentrations: out of the equilibrium of phase separation

We next consider the case when the timescales of gene expression and phase separation dynamics become comparable. To this end, we modify the gene network from (27) to account for phase separation by extending it by the stochastic exchange reactions from (24). The overall reaction system then becomes

$$\begin{aligned}
\emptyset &\xrightarrow{\Lambda_r} \mathrm{N}_r \xrightarrow{n_r/\tau_r} \emptyset \\
\emptyset &\xrightarrow{\lambda_p n_r} \mathrm{N}_+ \xrightarrow{n_+/\tau_p} \emptyset \\
&\mathrm{N}_+ \underset{k_{out}(n_+,n_-)}{\overset{k_{in}(n_+,n_-)}{\rightleftarrows}} \mathrm{N}_- \\
&\mathrm{N}_- \xrightarrow{n_-/\tau_p} \emptyset,
\end{aligned} \tag{33}$$

where we now distinguish between proteins in the bulk and droplet phase, denoted by $\mathrm{N}_+$ and $\mathrm{N}_-$, respectively. Our goal is to quantify the mean and noise strength of the droplet and bulk phase protein concentration. For a given set of parameters and a particular realization of the random transcription rate $\Lambda_r = \lambda_r$, the dynamics of system eq. (33) can be described by a master equation of the form

$$\begin{aligned}
\frac{\mathrm{d}}{\mathrm{d}t}\pi\left(n_r, n_+, n_-\right) =& \pi\left(n_r - 1, n_+, n_-\right)\lambda_r \\
&+ \pi\left(n_r + 1, n_+, n_-\right)\left(n_r + 1\right)\tau_r^{-1} \\
&+ \pi\left(n_r, n_+ - 1, n_-\right)\lambda_p n_r \\
&+ \left[\pi\left(n_r, n_+ + 1, n_-\right)\left(n_+ + 1\right) + \pi\left(n_r, n_+, n_- + 1\right)\left(n_- + 1\right)\right]\tau_p^{-1} \\
&+ \pi\left(n_r, n_+ + 1, n_- - 1\right)k_{in}\left(n_+ + 1, n_- - 1\right) \\
&+ \pi\left(n_r, n_+ - 1, n_- + 1\right)k_{out}\left(n_+ - 1, n_- + 1\right) \\
&- \pi\left(n_r, n_+, n_-\right)\big[\lambda_r + n_r\left(\tau_r^{-1} + \lambda_p\right) + \left(n_+ + n_-\right)\tau_p^{-1} \\
&\qquad\qquad + k_{in}\left(n_+, n_-\right) + k_{out}\left(n_+, n_-\right)\big],
\end{aligned} \tag{34}$$

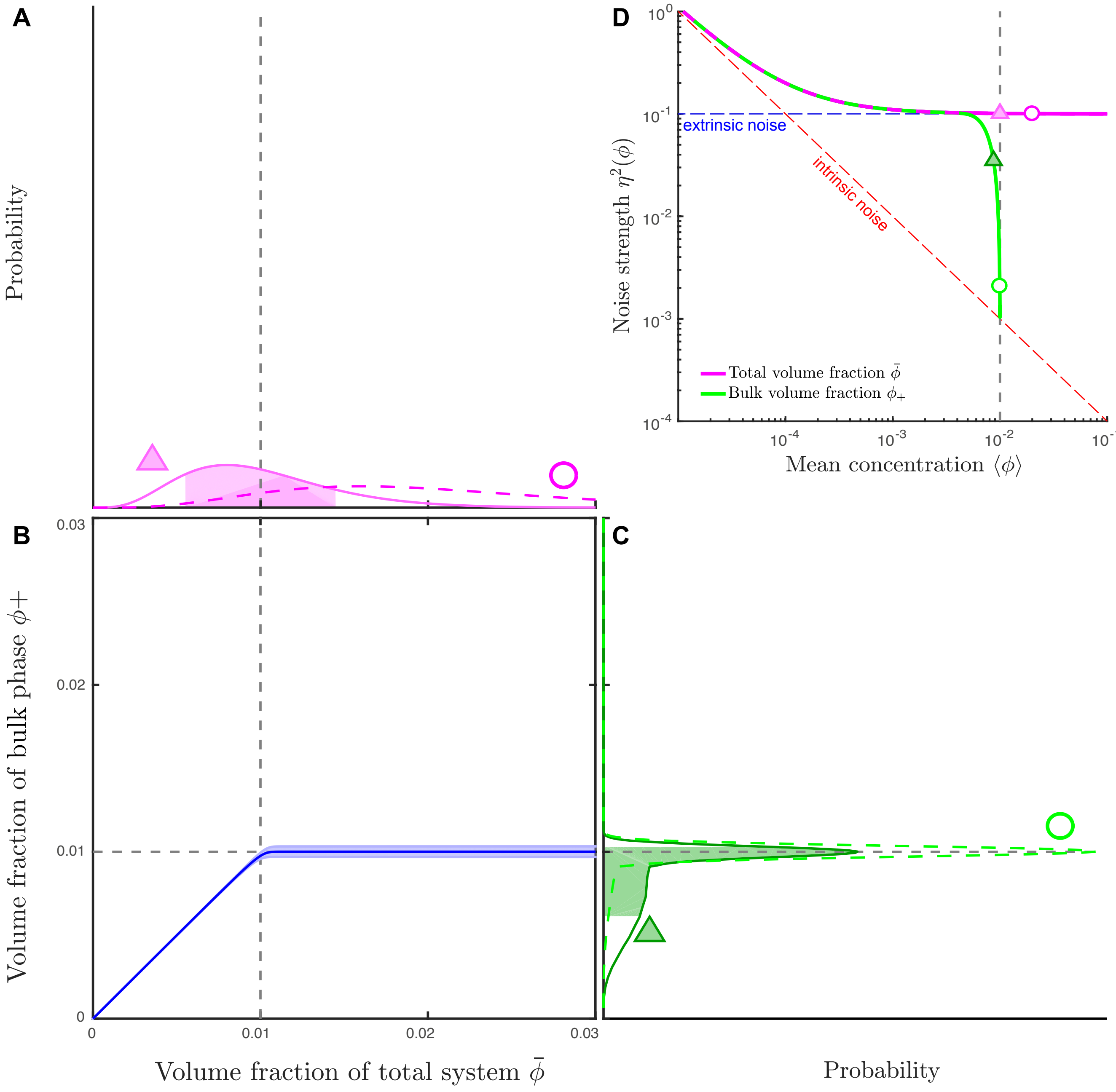

**Figure ST4:** Buffering in the limit of infinitely fast droplet dynamics. **A**: The negative binomial distribution for two different mean expression levels. The phase separation properties illustrated in **B** provide a conversion from the given total volume fractions to a buffered volume fractions of the bulk phase, which are shown in **C**. The resultion noise strenght for negative binomial distribution with varying parameter $p$, i.e. different mean expression levels, is shown in **D**. Grey dashed lines indicate the threshold concentration for phase separation in the thermodynamic limit (see Section 1.1.3). Shaded areas in A, B and C indicated the range of one standard deviation from mean in each direction of the respective distribtuion. The two noise characteristics of the two exemplary distributions shown in A and C are marked in D by triangles and circles, respectively. The red dashed line in D indicates the intrinsic noise limit and the blue dashed line the extrinsic noise limit, $\eta^2(\Lambda_r)$. The parameters are set to the same values as for Figure 3B and 3C in the main paper and are noted in Tab. 4 on page 14.

with $\pi(n_r, n_+, n_-) = \mathcal{P}(N_r(t) = n_r, N_+(t) = n_+, N_-(t) = n_- \mid \Lambda_r = \lambda_r)$. In order to compute the means and noise strengths, we first calculate the conditional moments

$$\langle \Phi_+ \mid \Lambda_r = \lambda_r \rangle = \sum_{n_-=0}^{\infty} \sum_{n_+=0}^{\infty} \sum_{n_r=0}^{\infty} \phi_+(n_+, n_+ + n_-)\pi(n_r, n_+, n_-) \tag{35}$$

$$\sigma^2(\Phi_+ \mid \Lambda_r = \lambda_r) = \sum_{n_-=0}^{\infty} \sum_{n_+=0}^{\infty} \sum_{n_r=0}^{\infty} \Big(\phi_+(n_+, n_+ + n_-) - \langle \Phi_+ \mid \Lambda_r = \lambda_r \rangle\Big)^2 \pi(n_r, n_+, n_-) \tag{36}$$

$$\big\langle \bar{\Phi} \mid \Lambda_r = \lambda_r \big\rangle = \sum_{n_-=0}^{\infty} \sum_{n_+=0}^{\infty} \sum_{n_r=0}^{\infty} \bar{\phi}(n_+ + n_-)\pi(n_r, n_+, n_-) \tag{37}$$

$$\sigma^2(\bar{\Phi} \mid \Lambda_r = \lambda_r) = \sum_{n_-=0}^{\infty} \sum_{n_+=0}^{\infty} \sum_{n_r=0}^{\infty} \Big(\bar{\phi}(n_+ + n_-) - \big\langle \bar{\Phi} \mid \Lambda_r = \lambda_r \big\rangle\Big)^2 \pi(n_r, n_+, n_-) \tag{38}$$

and subsequently average these moments over all possible realizations of $\lambda_r$, i.e.,

$$\langle \Phi_+ \rangle = \langle \langle \Phi_+ \mid \Lambda_r = \lambda_r \rangle \rangle \tag{39}$$

$$\sigma^2(\Phi_+) = \Big\langle \sigma^2(\Phi_+ \mid \Lambda_r = \lambda_r) + \langle \Phi_+ \mid \Lambda_r = \lambda_r \rangle^2 \Big\rangle - \langle \Phi_+ \rangle^2 \tag{40}$$

$$\big\langle \bar{\Phi} \big\rangle = \big\langle \big\langle \bar{\Phi} \mid \Lambda_r = \lambda_r \big\rangle \big\rangle \tag{41}$$

$$\sigma^2(\bar{\Phi}) = \Big\langle \sigma^2(\bar{\Phi} \mid \Lambda_r = \lambda_r) + \big\langle \bar{\Phi} \mid \Lambda_r = \lambda_r \big\rangle^2 \Big\rangle - \big\langle \bar{\Phi} \big\rangle^2 . \tag{42}$$

However, computation of the conditional moments (35-38) via direct numerical integration of (34) is computationally prohibitive, especially when RNAs and proteins are highly abundant. We, therefore, employ the linear noise approximation (LNA) [5, 6], which approximates $\pi(n_r, n_+, n_-)$ in terms of a Gaussian distribution

$$\pi(n_r, n_+, n_-) \approx N\left(m(t), \Sigma(t)\right), \tag{43}$$

with approximate moments

$$m(t) \approx \mathrm{E}\left[N(t) \mid \Lambda_r = \lambda_r\right]$$

and

$$\Sigma(t) \approx \mathrm{E}\left[(N(t) - m(t))(N(t) - m(t))^T \mid \Lambda_r = \lambda_r\right]$$

for $N(t) = (N_r(t), N_+(t), N_-(t))^T$. The moments $m(t)$ and $\Sigma(t)$ are described by ordinary differential equations obtained by considering the leading order terms of van Kampen's system size expansion applied to (34). Once $m(t)$ and $\Sigma(t)$ have been solved for, (35-38) can be efficiently evaluated via numerical integration. This provides an efficient way to compute the relationship between mean and noise strength of the total and bulk phase concentration when gene expression and phase separation have arbitrary time scales.

We finally remark that full probability distributions over the three species $N_r(t)$, $N_+(t)$ and $N_-(t)$ are straightforward to compute by integrating the Gaussian distribution from (43) with respect to $p(\lambda_r)$, for instance using quadrature methods. This will be required later in Section 2.3 where we compare the theoretical predictions with the experimental data.

### 2 Comparison between theory and experiment

In order to compare the theoretical model to experimental data, we require quantitative knowledge of the kinetic parameters of phase separation and protein expression. In Section 2.2, we will describe a Bayesian inference procedure, which we employed to calibrate our model on experimental single-cell data. Subsequently, we will discuss how we compared the model predictions to the experimental data.

#### 2.1 Experimental time-series data of 2NT-DDX4$^{\mathrm{YFP}}$ expression

As described in Materials and Methods, we performed experimental time-lapse experiments in which we tracked 2NT-DDX4$^{\mathrm{YFP}}$ expression in 93 cells over multiple hours post transfection. Measurements of 2NT-DDX4$^{\mathrm{YFP}}$ concentrations across the whole nucleus and the bulk phase were taken every ten minutes. Additionally, the measurements revealed cross sections of the entire nucleus and droplet phase. By considering both a spherical nucleus and a spherical droplet we estimated the corresponding volumes from the cross section areas. This multiplied by the measured concentration results in protein copy numbers for each phase, each cell and each time point.

The time-series were manually curated to exclude outlier cells. Moreover, the first eight time points were removed to ensure that all considered cells have exited mitosis and started expression. Additionally, we only considered time points where at least 30 data points were available. The complete data set is shown in Fig. ST5, where curated data points used for subsequent analyses are shown in green. From the resulting data, we computed the mean and variance of the total and bulk phase 2NT-DDX4$^{\mathrm{YFP}}$ copy numbers for each time point (Fig. 4 in the main text), which we used to determine the model parameters as described in the following Section 2.2.

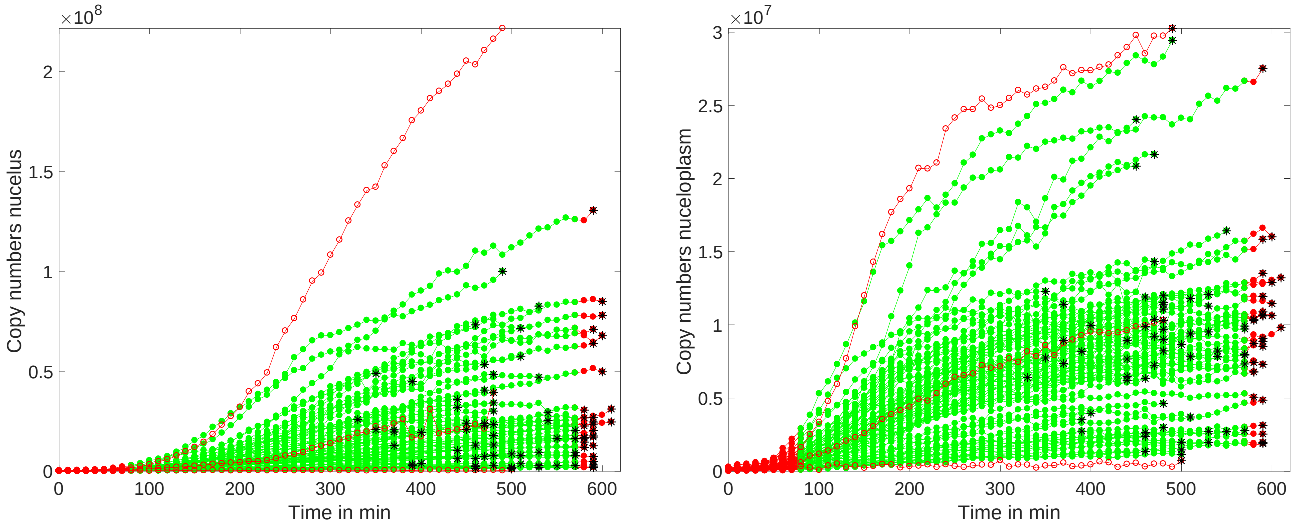

(a) Total amount

(b) Amount in bulk phase

**Figure ST5:** Copy numbers of 2NT-DDX4$^{\mathrm{YFP}}$ in the whole nucleus (a) and in the bulk phase of the nucleus (b) for the time-series experiment. Circles and disks mark each single measured value. Green indicates measurement point that where used for the Bayesian inference, while red points where not included. Black asterisks mark the end of the respective trajectories.

#### 2.2 Model calibration by Bayesian inference

In total, our model comprises several parameters associated with transcription and translation $(\langle\Lambda_r\rangle, \eta^2(\Lambda_r), \tau_r, \lambda_p, \tau_p)$ as well as phase separation $(\phi^*, T, V_{tot}, v, D)$. Some of these parameters, however, can be eliminated or determined directly from the experimental measurements. An estimate of the total system volume $V_{tot}$ was obtained from the data by taking the average nuclear volume across all time points and cells. The effective molecular volume $v$ was calculated as the average of the inverse 2NT-DDX4$^{\mathrm{YFP}}$ concentration inside the droplet phase (see also eq. (2)). Moreover, we found that the surface tension $\gamma$ has negligible influence on the dynamics of phase separation when set to realistic values. In particular, it was shown in [7] that the droplet surface tension *in vivo* was in the order of $1\mu\mathrm{N/m}$ in which case it had no noticeable effect on the dynamics of our model. We thus set $\gamma = 0$ to eliminate this parameter.

Furthermore, we can remark that the free energy enters the system dynamics only via the Boltzmann factor, such that the thermodynamic properties of phase separation are fully captured by the following term:

$$\begin{aligned}\frac{\tilde{F}\left(n_{+},\bar{n}\right)}{k_B T} &= \frac{F\left(n_{+},\bar{n}\right)}{k_B T} + \frac{\mu}{k_B T}\bar{n} \\ &= \frac{\mu}{k_B T}n_{+} + n_{+}\ln\left(\frac{n_{+}}{(V_{tot}/v - \bar{n} + n_{+})}\right) - n_{+} \\ &= \left[\phi^{*} - \ln\left(\phi^{*}\right) + \ln\left(\frac{n_{+}}{(V_{tot}/v - \bar{n} + n_{+})}\right) - 1\right] n_{+},\end{aligned} \tag{44}$$

where we have used eq. (14) in the last step. Thus, we are left with only one free parameters that characterizes the equilibrium thermodynamic properties of phase separation, i.e., the threshold volume fraction $\phi^{*}$. To fix the threshold volume fraction $\phi^{*}$ we took the concentration of the bulk phase for the first time point of each trajectory, which exhibited protein droplets and average this value over all trajectories which showed phase separation. In total, this we are left with six unknown parameters, describing the dynamics of the system, as well as two additional fit parameters corresponding to the mean and variance of the initial mRNA abundance.

In order to quantitate these parameters from the 2NT-DDX4$^{\mathrm{YFP}}$ time-series experiments, we used a Bayesian inference approach. In particular, we resorted to an efficient moment-based inference scheme [8], which is based on matching the time course of the experimental moments (e.g., means and variances) with those obtained from the model using a Markov chain Monte Carlo (MCMC) technique. For more information on this method, the reader should refer to [8]. We performed the inference scheme on our data using lognormal proposal distributions for $M = 10000$ iterations and extracted maximum a posterior (MAP) estimates from it. This procedure was repeated eight times with randomly drawn initial conditions from which we selected the parameter set with the highest posterior probability. The resulting parameter values are shown in Table 1 and Table 2. We remark that in our model, the considered reaction volume corresponds to the nuclear volume instead of the total cell volume. Therefore, the inferred protein translation and degradation rates have to be understood as effective rates, subsuming nuclear import and export, respectively. The relatively small protein half life ($\tau_p = 1.2h$), for instance, is likely due to fast nuclear export since YFP typically degrades on the time scale of several hours.

| parameter | $V_{tot}$ | $v$ | $\phi^{*}$ | $\phi^{*}/v$ |
|---|---|---|---|---|
| value | $2.1 \cdot 10^{3}$ $\mu$m$^{3}$ | $1.5 \cdot 10^{3}$ nm$^{3}$ | $4.9 \cdot 10^{-3}$ | 5.5 $\mu$M |

**Table 1:** Directly measured model parameter

| parameter | $D$ | $\langle\lambda_r\rangle$ | $\eta^{2}\left(\lambda_r\right)$ | $\tau_r$ | $\lambda_p$ | $\tau_p$ | $\langle N_r(t = 80\text{min})\rangle$ | $\sigma(N_r(t = 80\text{min}))$ |
|---|---|---|---|---|---|---|---|---|
| value | 0.034 $\mu$m$^{2}$/s | 15 min$^{-1}$ | 1.3 | 7.2 h | 60 min$^{-1}$ | 1.2 h | $3.1 \cdot 10^{2}$ | $4.5 \cdot 10^{2}$ |

**Table 2:** Inferred model parameter

#### 2.3 Determining the relationship between mean and noise strength from experimental data

In order to compare between theory and experiment, we determined relationship between mean and noise strength of the bulk phase 2NT-DDX4$^{\mathrm{YFP}}$ concentration from our high-throughput single-cell dataset (Materials and Methods). To achieve this, we solved for the stationary distributions over protein copy numbers $\mathcal{P}(n_r, n_+, n_-)$ of system (33) for a fixed $\eta^{2}(\Lambda_r) = 0.15$ and different $\langle\Lambda_r\rangle$. For each value of $\langle\Lambda_r\rangle$, we generated corresponding subpopulations from the experimental data such that the distribution over the total nuclear 2NT-DDX4$^{\mathrm{YFP}}$ abundance was consistent with the theoretically obtained marginal distribution $\mathcal{P}(\bar{N} = \bar{n})$. To this end, we drew $5 \cdot 10^{3}$ random transcription rates for a given $\langle\Lambda_r\rangle$ and $\eta^{2}(\Lambda_r)$ and for each of them computed the stationary distribution over total protein copy numbers $\bar{N}$, which when using the LNA, is given by a Gaussian distribution. For each transcription rate, we then sampled $2 \cdot 10^{2}$ values of $\bar{N}$ and converted them to volume fractions $\bar{\Phi}$. In total, this gave rise to $10^{6}$ different samples of $\bar{\Phi}$ and to each of them, we assigned the experimental measurement with the closest total 2NT-DDX4$^{\mathrm{YFP}}$ volume fraction. Importantly, this producedure ensures that the theoretical and experimental subpopulations are consistent with each other in terms of the total volume fraction $\bar{\Phi}$ (see Figure ST6). Each of the experimental

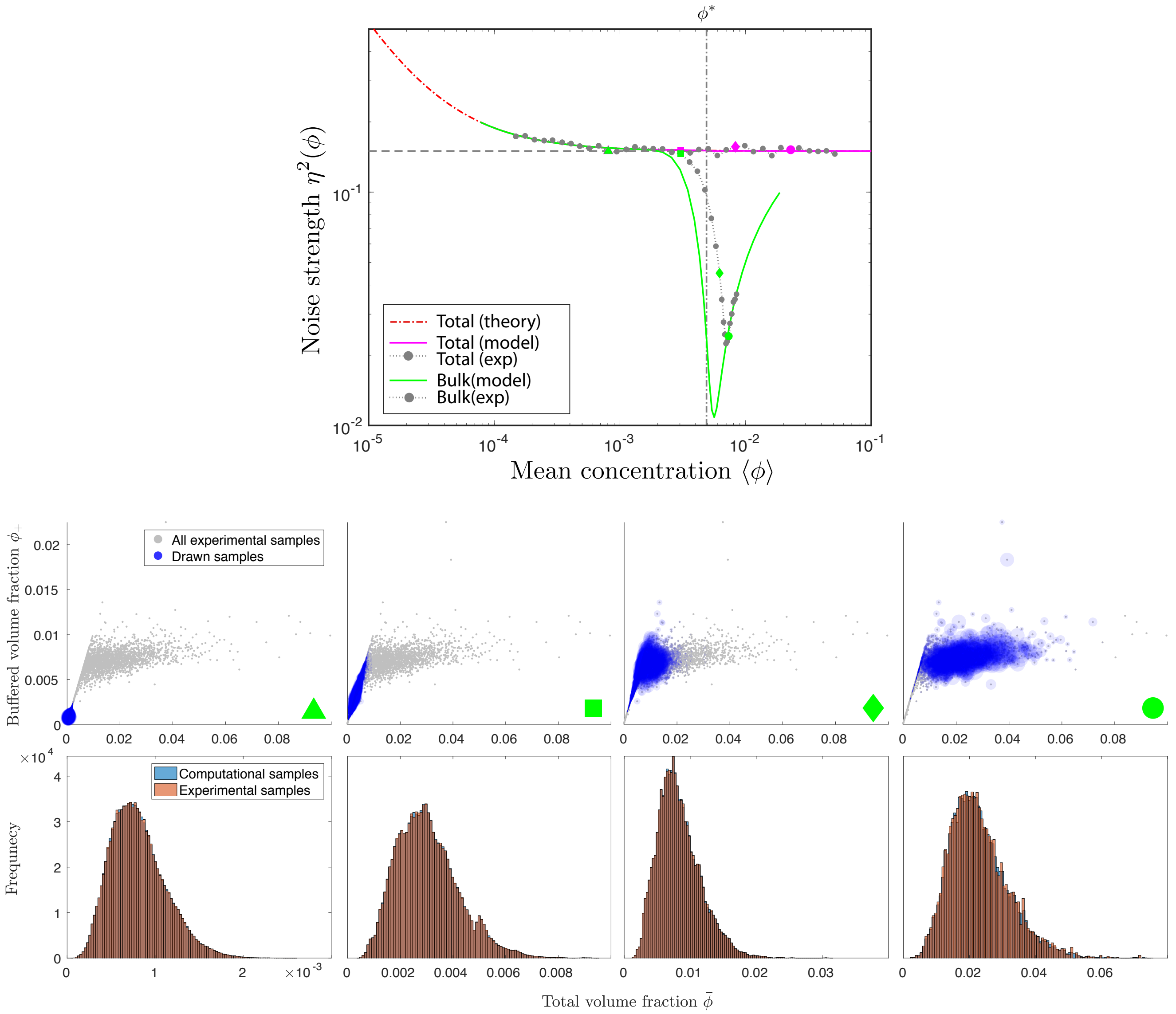

**Figure ST6:** Sampling subpopulations from the experimental single-cell data. The data used for the top plot is the same as for Fig 4E of the main text. The sampling procedure is illustrated for four data points indicated by the green and magenta triangles, squares, diamonds and circles. The middle panel shows scatter plots of all single-cell measurements of $\bar{\phi}$ and $\phi_+$ (grey). The blue dots indicate the subsampled measurements corresponding to four particular values of $\langle\bar{\phi}\rangle$ and $\eta^2(\bar{\phi})$ along the mean-noise curve as indicated with a triangle, square, diamond and circle, respectively. The size of each dot indicates how often the respective sample was drawn. The bottom panel shows that the resulting histograms over the samples of $\bar{\phi}$ generated from the model and drawn from the experimental data are consistent.

samples is associated with a corresponding measurement of the bulk phase volume fraction, for which we computed averages and noise strengths. In summary, this allowed us to determine the relationship between mean and noise strength of the bulk phase concentration, which we compared to the theoretically predicted one (Figure 4 in the main text).

#### 3 Parameter values used for the figures in the main paper

Here we list all the parameter values as they were used for Figure 2 and Figure 3 in the main text.

| parameter | $v$ | $V_{tot}/v$ | $T$ | $\phi^*$ | $\gamma$ |
|---|---|---|---|---|---|
| value | $10^{-25}$ m$^3$ | $10^4$ | 305 K | $10^{-2}$ | $10^{-6}$ N/m |

**Table 3:** Model parameters for Figure 2 in their respective SI units.

| parameter | $v$ | $V_{tot}/v$ | $T$ | $\phi^*$ | $\gamma$ | $\eta^2(\Lambda_r)$ |
|---|---|---|---|---|---|---|
| value | $10^{-25}$m$^3$ | $10^5$ | 305 K | $10^{-2}$ | $10^{-6}$N/m | 0.1 |

**Table 4:** Model parameters for Figure 3B & 3C and S4 in their respective SI units.

| parameter | $v$ | $V_{tot}/v$ | $T$ | $\phi^*$ | $\gamma$ | $\tau_r$ | $\lambda_p$ | $\tau_p$ |
|---|---|---|---|---|---|---|---|---|
| value | $10^{-25}$ m$^3$ | $10^5$ | 305 K | $10^{-2}$ | 0 | 69.31 | $10^{-2}$ | $6.931 \cdot 10^3$ |

**Table 5:** Model parameters for Figure 3D & 3D in their respective SI units.

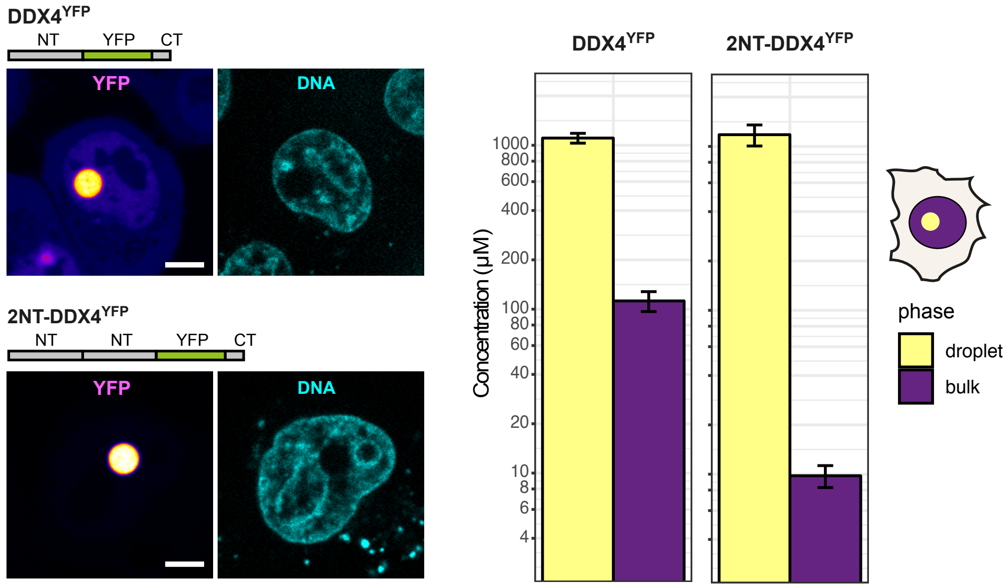

**Figure S1. Duplication of the N-terminus in DDX4$^{YFP}$ promotes phase separation by reducing saturation concentration.** Quantification of the droplet phase and the bulk nucleoplasmic phase in transfected HeLa cells containing large droplets formed by DDX4$^{YFP}$ (n=15 cells) or 2NT-DDX4$^{YFP}$ (n= 25 cells). Concentration was estimated based on mean fluorescence intensity and using a calibration curve made with serial dilutions of purified DDX4$^{YFP}$ protein. Images were taken 24 hours after transfection using 100x objective. The DNA was stained with 200 ng/ml of Hoechst dye. Scale bar = 10 $\mu$m.

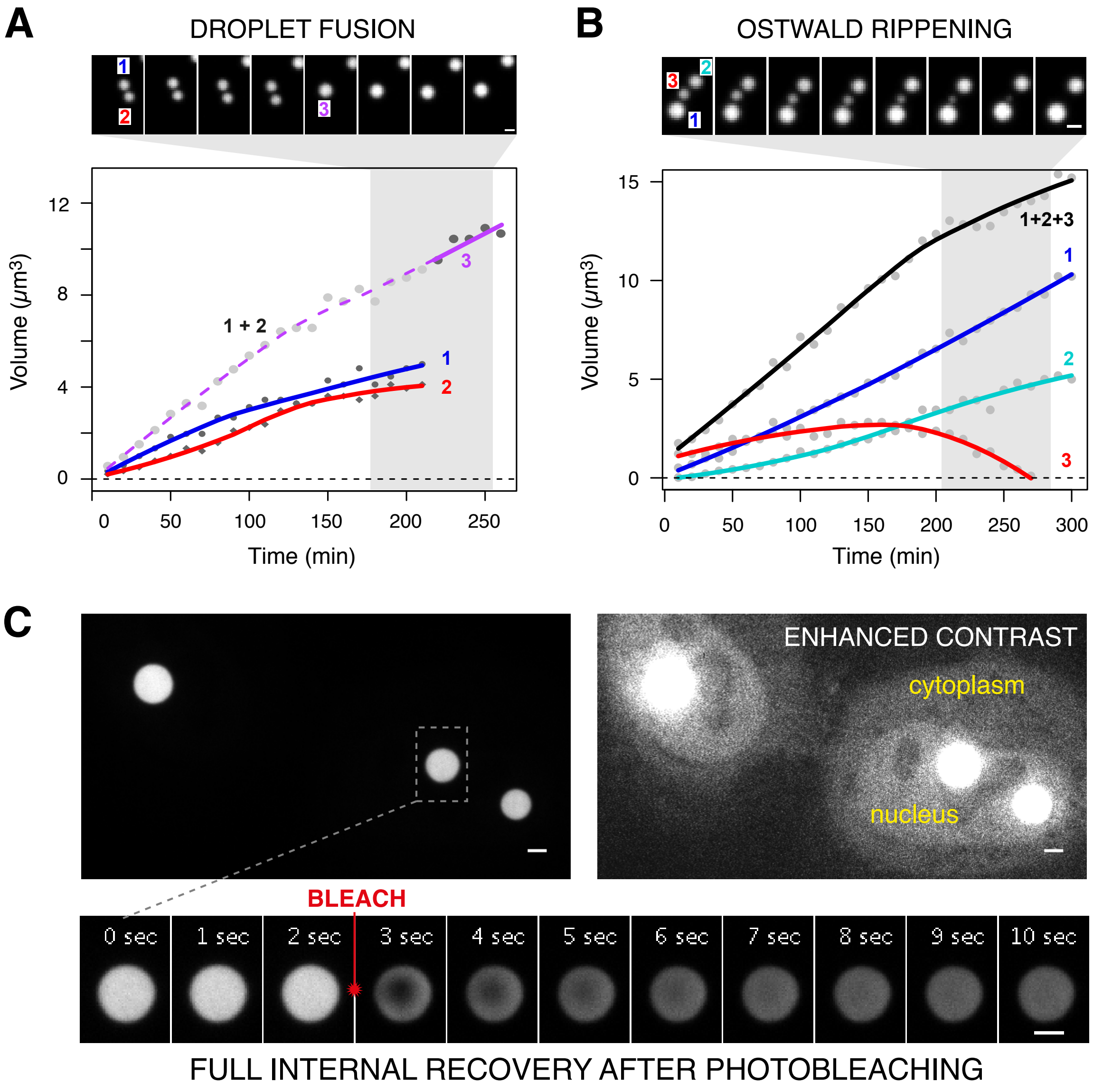

**Figure S2. Nuclear compartments formed by 2NT-DDX4$^{YFP}$ have liquid-like properties.** Growth of 2NT-DDX4$^{YFP}$ droplets through fusion (a) and Ostwald ripening (b) quantified by time-lapse microscopy and measurement of droplet volume over time. Local polynomial regression line (colored lines) was fitted through each data set. (c) 2NT-DDX4$^{YFP}$ droplets show fast and complete internal recovery upon photobleaching. The protein is enriched in the nucleus (nuc) as evident in the 'enhanced contrast' image and forms droplets only in that compartment with no evidence for condensate formation in the cytoplasm (cyt). Scale bar = 2 $\mu$m.

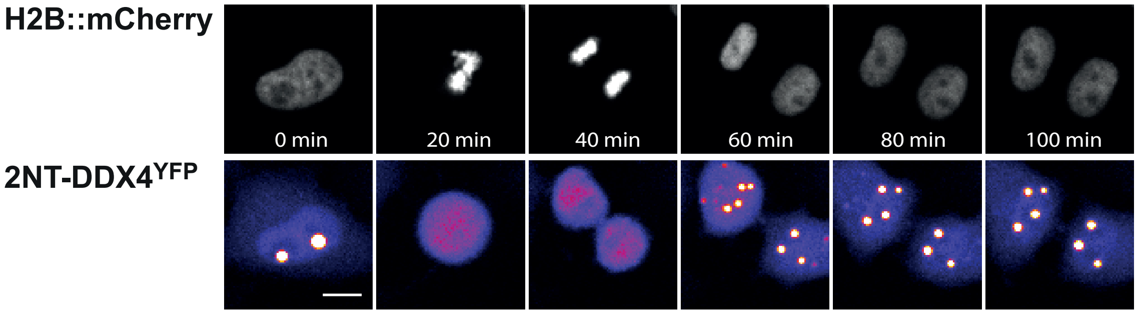

**Figure S3. 2NT-DDX4$^{YFP}$ droplets dissolve during mitosis.** Time-lapse imaging of a HeLa cell undergoing mitosis. Histone protein H2B tagged with mCherry expressed from a stable plasmid was as a marker for the chromatin and facilitated identification of mitotic cells. Scale bar = 10 $\mu$m.
